# Genome-wide association studies reveal a rapidly evolving candidate avirulence effector in the Cercospora leaf spot pathogen *Cercospora beticola*

**DOI:** 10.1101/2023.06.11.544483

**Authors:** Chen Chen, Harald Keunecke, Felix Bemm, Gabor Gyetvai, Enzo Neu, Friedrich J. Kopisch-Obuch, Bruce A. McDonald, Jessica Stapley

## Abstract

The major resistance gene *BvCR4* recently bred into sugar beets provides a high level of resistance to Cercospora leaf spot caused by the fungal pathogen *Cercospora beticola*. The occurrence of pathogen strains virulent to *BvCR4* was studied using field trials in Switzerland. Virulence of a subset of these strains was evaluated in a field trial conducted under elevated artificial disease pressure. We created a new *C. beticola* reference genome and mapped whole genome sequences of 256 field-collected isolates. These were combined with virulence phenotypes to conduct three separate GWAS to identify candidate avirulence genes. We identified a locus associated with avirulence containing a single candidate avirulence effector gene named *AvrCR4*. All virulent isolates either lacked *AvrCR4* or had non-synonymous mutations within the gene. *AvrCR4* was present in all 74 isolates from non-*BvCR4* hybrids, whereas 33 of 89 isolates from *BvCR4* hybrids carried a deletion. We also mapped genomic data from 190 publicly available U.S. isolates to our new reference genome. The *AvrCR4* deletion was found in only one of 95 unique isolates from non-*BvCR4* hybrids in the U.S. *AvrCR4* presents a unique example of an avirulence effector in which virulent alleles have only recently emerged. Most likely these were selected out of standing genetic variation after deployment of *BvCR4*. Identification of *AvrCR4* will enable real-time screening of *C. beticola* populations for the emergence and spread of virulent isolates.

## Introduction

Plant pathogens pose significant risks to crop production and food security around the world (Savary et al., 2019). Disease resistance genes (R-genes) provide economical and ecologically friendly approaches to disease control that are important for the production of many crops. Plant breeders often select for R-genes with major effects that form a critical part of the plant’s immune system. Major R-genes typically encode immune receptors, e.g. receptor-like kinases (RLKs) or nucleotide-binding leucine-rich repeat proteins (NB-LRRs) that detect pathogen proteins and trigger an immune response (Deng et al., 2020; Petit-Houdenot and Fudal, 2017). In sugar beet (*Beta vulgaris ssp. vulgaris*), R-genes have provided the only effective control for beet cyst nematode (*Heterodera schachtii*) (Stevanato et al., 2015) and rhizomania (Beet necrotic yellow vein virus) (Capistrano-Gossmann et al., 2017).

The effectiveness of R-genes can be reduced when virulent strains of the pathogen evolve. This evolution can happen within a few years, with well-documented cases including 2-7 years for *Leptosphaeria maculans* on oilseed rape (Rouxel and Balesdent, 2017; Van De Wouw et al., 2010); 3 years for *Zymoseptoria tritici* on wheat (Cowger et al., 2000), and 2-3 years for *Pyricularia oryzae* on rice (Kiyosawa, 1982). However, even when the pathogen population contains virulent strains, R-genes can still prove useful, e.g. the *Rlm7* gene encoding resistance to *L. maculans* remains useful more than 15 years after deployment despite the identification of virulent isolates (Alnajar et al., 2022; Winter and Koopmann, 2016). Understanding the mechanisms governing the emergence of virulence is needed to develop more effective strategies to mitigate the impact of pathogens in agro-ecosystems. Plant-pathogen coevolution in natural ecosystems created the reservoir of R-genes that can be integrated into domesticated crops via plant breeding programs. These R-genes typically recognise pathogen molecules, often called effectors, that function to increase susceptibility of the host to pathogen infection (Kamoun, 2007). If the recognition of an effector by an R-protein leads to an immune response that severely limits pathogen infection, it is called an avirulence effector. The genes encoding avirulence effectors typically encode small, secreted, cysteine-rich proteins with limited homology to other characterized proteins (Rouxel and Balesdent, 2017; Sonah et al., 2016). Many avirulence effector genes have now been identified and cloned in several important plant pathosystems, including *L. maculans* on oilseed rape (Rouxel and Balesdent, 2017), *P. oryzae* on rice (Fernandez, 2023), and *Z. tritici* on wheat (Meile et al., 2018; Zhong et al., 2017).

Following deployment of an R-gene, the corresponding avirulence effector is under strong directional selection and typically evolves rapidly to evade host recognition, creating virulent pathogen strains that can infect plants carrying the R-gene. This evolution can involve gene deletions, non-synonymous mutations, and alterations in gene transcription. The genes encoding fungal effectors are often located in dynamic, TE-rich regions of the genome that facilitate an accelerated evolution for these genes (Lo Presti et al., 2015). After years of selection imposed by a widely deployed R-gene, the gene encoding the corresponding avirulence effector typically displays very high levels of genetic diversity, while the original wild-type sequence of the effector gene may have disappeared entirely or be present at only very low frequencies (Brunner et al., 2007; Brunner and McDonald, 2018; Dodds et al., 2006; Stephens et al., 2021).

Sugar beet provides around one-fifth of global sucrose production every year (Rangel et al., 2020). Cercospora leaf spot (CLS), caused by the fungus *Cercospora beticola*, is the most common and destructive disease of sugar beet. Sugar beet cultivars with improved resistance to CLS have been under development for thirty years (Rangel et al., 2020). Wild sea beet (*Beta vulgaris* subsp. *maritima*) is the most important source of resistant genes and provided at least four quantitative trait loci (QTL) for resistance to CLS . It has proved difficult to combine high resistance and high yield into the same CLS-resistant cultivar (Setiawan et al., 2000; Taguchi et al., 2011). The most widely planted cultivars combine high yield and moderate CLS resistance (Windels et al., 1998), but rely mainly on fungicides to limit losses associated with CLS. Recently, Törjék et al (2022; 2020) identified an NBS-LRR resistance gene named *BvCR4* that showed a high level of CLS resistance when bred into sugar beets, enabling a unique combination of high CLS resistance and high yield in the same variety. However, *BvCR4* might be vulnerable to the evolution of virulent *C. beticola* strains if it were deployed in the absence of supporting quantitative background resistance (Pilet-Nayel et al., 2017) and without supplemental plant protection measures.

The evolutionary potential of *C. beticola* is high because it has a mixed reproductive system, exhibits extensive gene flow, and undergoes several cycles of reproduction during each growing season. Field populations contain high levels of genetic diversity as expected for pathogen populations that maintain a high effective population size (Vaghefi et al., 2017). *C. beticola* has already shown the capacity to rapidly evolve resistance to several fungicides representing different modes of action, including triazole, benzimidazole, difenoconazole, organotin, sterol demethylation inhibitor (DMI), and quinone outside inhibitor (QoI) (Bolton et al., 2012; Karaoglanidis et al., 2000). This evolution of fungicide resistance underlines the need to make more effective use of CLS resistance, but also illustrates the potential for pathogen adaptation following the deployment of R-genes.

We conducted a series of field, greenhouse, and lab experiments to assess the risks associated with deployment of *BvCR4*. We monitored CLS epidemics in experimental hybrids carrying *BvCR4* and isolated putative virulent strains of *C. beticola* from infected leaves. We confirmed the virulence of many of these strains on *BvCR4* hybrids growing under field conditions. We created a new reference genome sequence for *C. beticola*, analysed genome sequences for 256 isolates and conducted three GWAS analyses to search for candidate avirulence effectors. Each GWAS identified the same candidate avirulence effector (*AvrCR4*) on chromosome 1. The *AvrCR4* gene contained several different mutations encoding virulence and was deleted in many isolates coming from *BvCR4* hybrids.

## Results

We monitored CLS epidemics at two sites in Switzerland planted with four hybrids: two resistant hybrids carrying *BvCR4* and two susceptible hybrids without *BvCR4* (Table S1, Figure S1). Plots planted to resistant *BvCR4* hybrids showed later onset of CLS and slower epidemic progress compared to plots planted to susceptible hybrids. Total levels of CLS infection in the resistant hybrids never reached the levels found in the susceptible hybrids. Leaf death and subsequent regrowth was less prevalent in the resistant hybrids carrying *BvCR4*. Leaves with CLS lesions were collected throughout the season. Similar experimental plots were established in Germany, but diseased leaves were collected only at the end of the season (Figure S1). We made pure cultures of *C. beticola* and sequenced these isolates using Illumina short read sequencing (sample information see Table S2). Interestingly, one of the genotypes observed only on resistant hybrids carrying *BvCR4* was frequent and widely distributed at the Rudolfingen site, suggesting it was a virulent clone.

To assess the virulence of isolates we performed a field trial in 2022 in Northeim, Germany. Four sugar beet hybrids (Table 1) were inoculated, and the disease was scored using a 9- point rating scale (Table S3) at 32-, 40-, 47-, 56- and 64-days post inoculation (dpi) (Table S4). Fifty-two unique isolates from the Swiss plots were phenotyped in the virulence trial, 49 from plots planted with *BvCR4* hybrids (18 isolates from Hybrid_R2, 31 isolates from Hybrid_R3), and three were from susceptible hybrids (2 from Hybrid_S3, 1 from Hybrid_S4). We also included 5 separate isolations of the putatively virulent clone from the Rudolfingen site to test reproducibility of the phenotypes, but only one was used for GWAS analyses. Four additional reference inocula with known virulence properties were used as controls.

**Table 1.** Sugar beet hybrids used for the virulence assessments.

| Experimental Hybrids | *Cercospora* resistance level |
| --- | --- |
| Hybrid_S4 | low quantitative resistance |
| Hybrid_S3 | moderate quantitative resistance |
| Hybrid_R2 | *BvCR4* with low quantitative background resistance |
| Hybrid_R3 | *BvCR4* with moderate quantitative background resistance |

Disease ratings increased over the growing season, peaking at 56 dpi (Figure 1). Disease ratings at 32-56 dpi were highly correlated while the rating at 64 dpi was less correlated with other time points (Figure S2). Using the 40-dpi rating on Hybrid_R2, isolates were scored as virulent (>4) or avirulent (≤4), resulting in 25 of the 52 unique field-collected isolates scored as virulent and 27 scored as avirulent. The three isolates from susceptible hybrids were all avirulent. Next, we considered how dpi, hybrid, and virulence score (0/1) explained variation in the disease rating using linear models. We found significant pairwise interactions but no three-way interaction (Table S5). Disease ratings increased over time on all hybrids, but the pattern differed between hybrids with and without *BvCR4* (Figure 1). Isolates scored as virulent caused more disease at all timepoints on Hybrid_R3 carrying *BvCR4* compared to the avirulent isolates (it was not statistically appropriate to do this analysis on Hybrid_R2 because this was used to define virulent/avirulent). When tested on hybrids without *BvCR4* (Hybrid_S3 and Hybrid_S4), virulent and avirulent isolates caused similar levels of disease.

**Figure 1.**
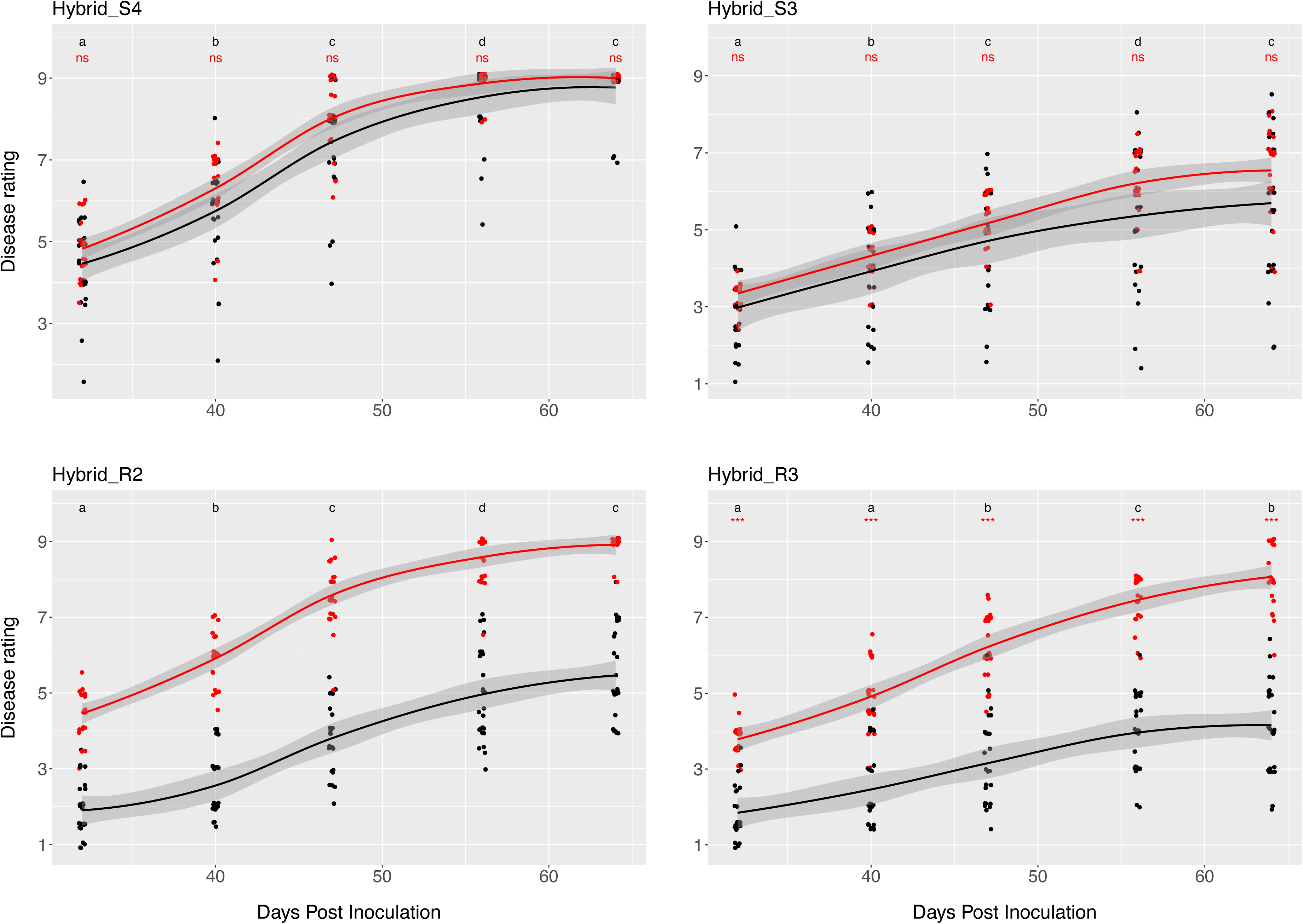
Disease rating of isolates at five time points in the field trial. Each panel shows the rating on one of the four hybrids (2 susceptible and 2 resistant). Red points are isolates scored as virulent (rating >4 on Hybrid_R2 at 40 dpi) and black are avirulent (rating ≤4 on Hybrid_R2 at 40 dpi), lines are loess-smoothed lines. Black letters at the top of the plots indicate least significant differences between mean rating of all isolates at each time point. Red text indicate difference in disease rating between virulent and avirulent for that hybrid and time point (ns=nonsignificant, *p<0.05, **p<0.01, *** p<0.001). No analysis was performed on Hybrid_K2 because data from this hybrid was used to score virulence/avirulence.

### Genome-wide association scans identified a single effector gene candidate

We performed three GWAS analyses; two based on SNPs and one based on k-mers. For the SNP-based GWAS, the sequence data was mapped to a high-quality reference genome that we created using Nanopore long read sequencing. The reference genome was assembled into 11 continuous sequences (hereafter called chromosomes). The genome showed a completeness of 98.4% according to the BUSCO benchmark against Ascomycota single copy orthologs, with 13,869 annotated genes (Table S6). We retained 358,365 high quality variants for the SNP GWAS. The first GWAS used the quantitative virulence rating on the resistant hybrid (Hybrid_R2) at 40 dpi as the phenotypic data (n=52), referred to hereafter as the ‘quantitative GWAS’. The resulting Manhattan plots showed a single distinctive, but not statistically significant, LOD peak on chromosome 1 (Figure 2a). The second GWAS coded the phenotype data as either virulent (Hybrid_R2 rating at 40 dpi > 4) or avirulent (Hybrid_R2 rating at 40 dpi ≤ 4) and included 38 additional isolates that were sampled from the susceptible hybrid Hybrid_S2 and scored as avirulent, giving a total sample size of 90. This GWAS identified a significant LOD peak on chromosome 1 with an interval spanning 6730 bp (4087634-4094364 bp) (Figure 2b). This peak was in the same position as the peak observed in the quantitative GWAS. The GWAS interval contained a single gene (ID=Hevensen_hq-v2.G01679) that is 340 bp long (Figure 3). The gene is predicted to encode an effector protein that is 89 amino acids (aa) long. The putative effector contains a signal peptide of 16aa (predicted by TOPCONS, Tsirigos et al., 2015) and is cysteine-rich (6%). We did not find a significant match using a BLAST search against the publicly available reference genome, but a partial hit was found on Chr 7. A copy of the gene with 88% identity was found in *Cercospora sojina* (strain RACE15) chromosome XII: 4346561-4346898.

**Figure 2.**
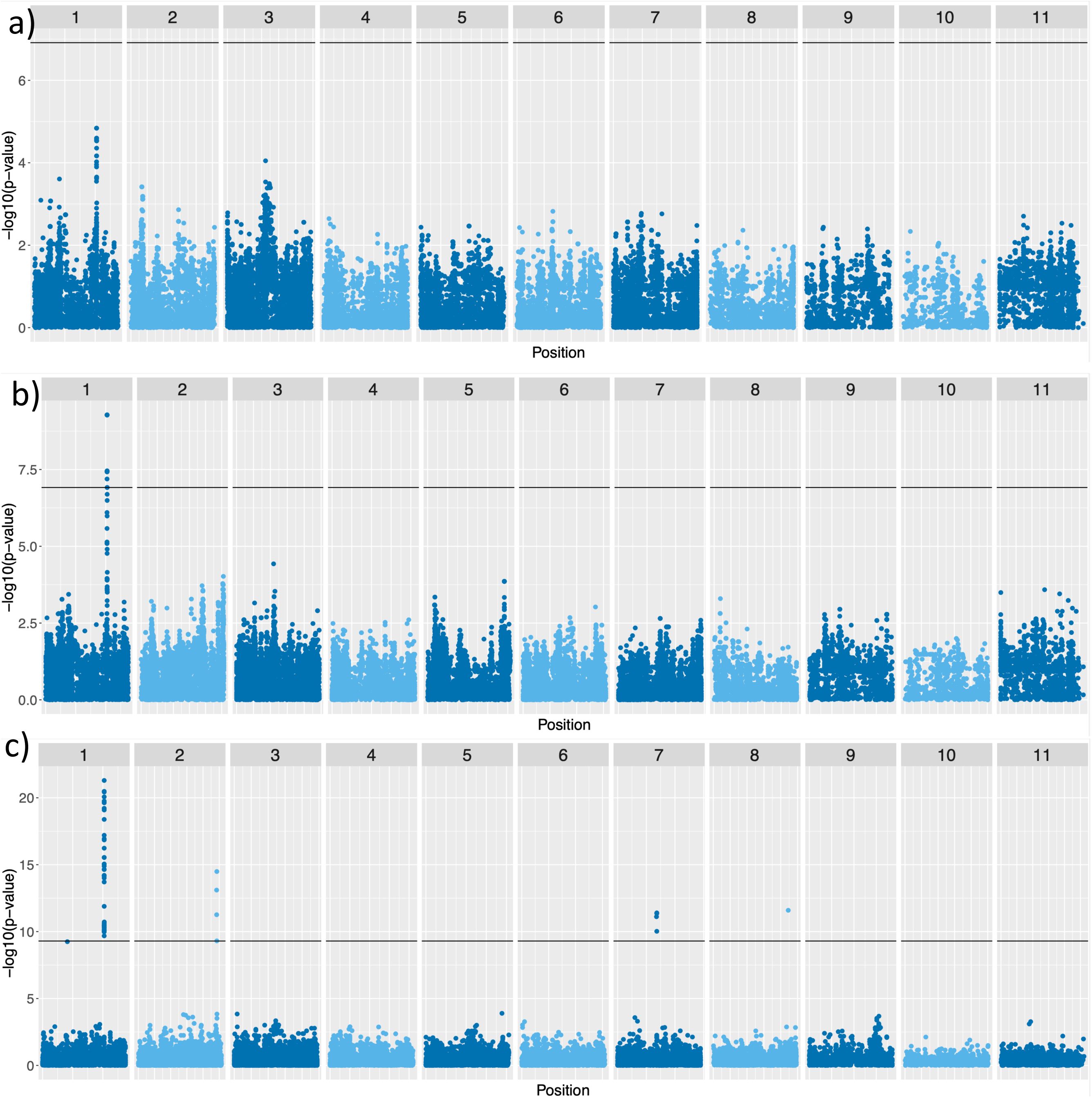
Manhattan plots of GWAS. a) Manhattan plot from a genome-wide association (GWAS) scan using 52 unique isolates with quantitative phenotypes at 40 dpi on Hybrid_R2. b) Manhattan plot from GWAS using 90 unique isolates, with 52 isolates scored with binary phenotypes (virulent or avirulent) and 38 isolates collected from Hybrid_S4 scored as avirulent. c) Manhattan plot from a k-mer GWAS using 47 unique isolates with quantitative phenotypes. Horizontal line is the genome-wide significance level. All SNPs and k-mers with p-value >1.22e-01 were downsampled for plotting.

**Figure 3.**
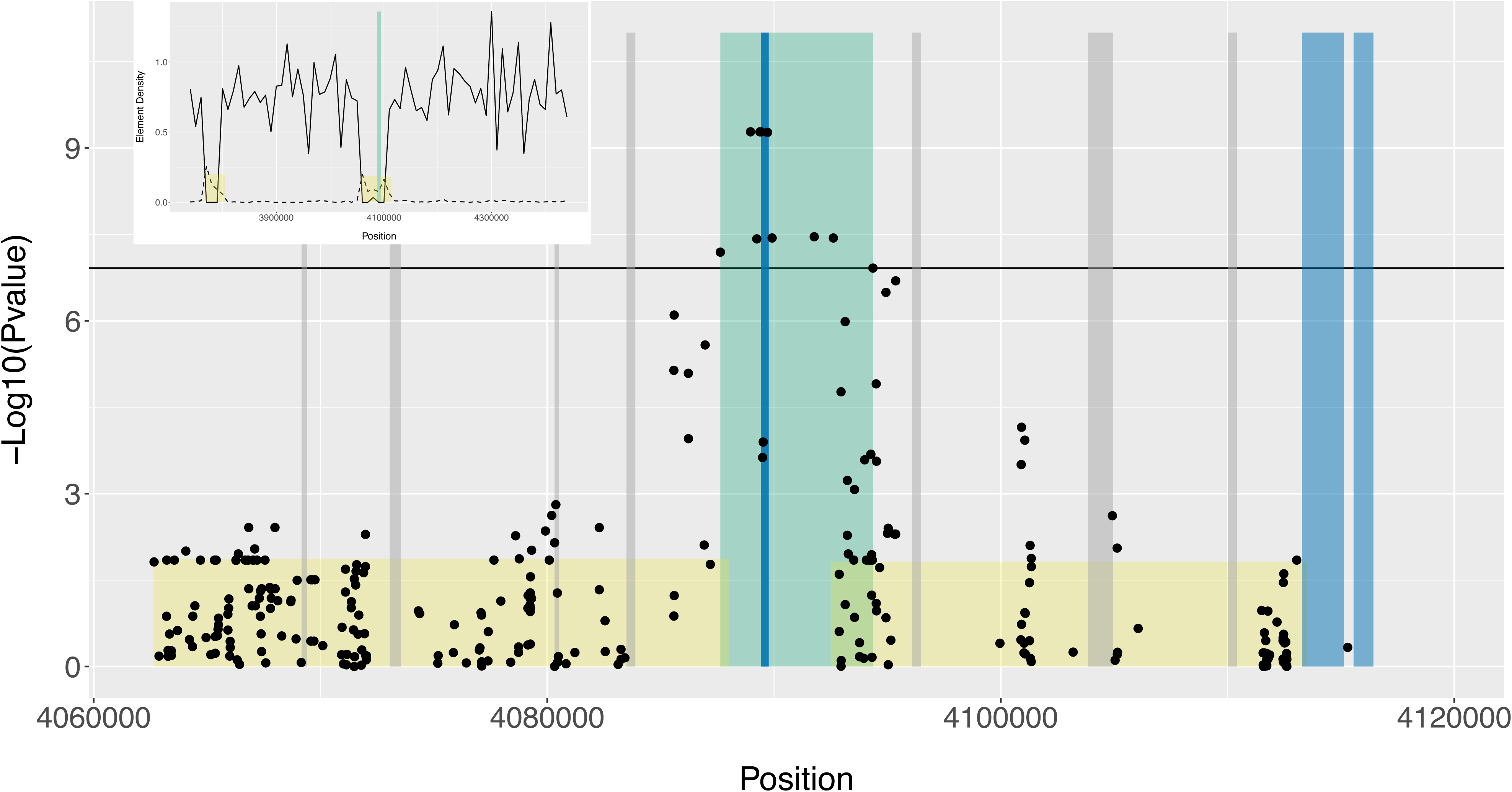
Manhattan plot of the significant GWAS interval (± 25 kb of flanking sequence) on chromosome 1:4087634-4094364 bp (shaded green). Genes are shaded in blue, full length transposable elements are shaded in grey and large RIP affected regions (LRAR) are shaded in yellow. Inset: Gene and repeat density around the GWAS interval (including 300 kb of flanking sequence). Gene (solid line) and repeat (dashed line) density shown in 10 kb windows. The GWAS interval is shaded in green and the LRAR is shaded in yellow.

The k-mer GWAS used the quantitative virulence rating on the resistant hybrid (Hybrid_R2) at 40 dpi (n=47). Several significant peaks were identified (Figure 2c). A peak on Chromosome 1 had significant k-mers located within the same interval identified in the previous analyses. Other significant associations containing more than one k-mer were found on Chromosomes 2 and 7. In both cases the k-mers with significant associations were not tightly clustered, unlike the peak on Chr 1. So, we consider these to be false positives. Significant singleton k-mers found on Chromosomes 1 (1644617bp) and Chromosome 8 (2223033bp) were also treated as false positives.

The effector gene candidate was in a gene-sparse, repeat-rich region of the genome that exhibits signatures of Repeat Induced Point mutations (RIP) (Figure 3). To investigate variation in the effector gene, we used genomic data from field collected isolates and publicly available sequence data from the Short Read Archive (SRA). After mapping, quality checking and removal of sequence duplicates and clones we retained 258 unique genomes (163 sequenced by us, 95 from SRA) for further analyses. Of these 258 unique genotypes, 34 had the candidate effector deletion. Among isolates collected from Switzerland and Germany, the gene was absent in over a third of isolates collected from resistant hybrids carrying *BvCR4* (37%) and was present in all isolates collected from susceptible hybrids (Table 2). In the publicly available genome sequences from the USA, the gene deletion was observed only once out of 95 unique genotypes (1%).

**Table 2.** Frequency of unique isolates carrying a deletion of the candidate avirulence effector gene. Frequency is the percentage of the total number of individuals analyzed. The number of individuals with deletions and the total are shown in parentheses.

| Country | Sampling | Year | Resistant:<br>Deletion freq. (n) | Susceptible:<br>Deletion freq. (n) | Clone<br>Corrected<br>(Total) |
| --- | --- | --- | --- | --- | --- |
| Switzerland | Pilot | 2019 | 25% (1/4) | NA | 4 (7) |
| Switzerland | Temporal | 2020 | 32% (22/68) | 0% (0/42) | 110 (147) |
| Switzerland | Transect | 2020 | 57% (4/7) | 0% (0/15) | 22 (38) |
| Germany | Transect | 2020 | 60% (6/10) | 0% (0/17) | 27 (40) |
| USA | Public | 2016-17 | NA | 1.0% (1/95) | 95 (190) |
| Total |  |  | 37% (33/89) | 0.5% (1/169) | 258 (422) |

### Sequence variation in the candidate effector gene

Among the 224 isolates carrying the candidate effector gene, 195 (87%) had no variation in the gene (158 from susceptible hybrids, 37 from resistant hybrids). Among the remaining 29 isolates, we identified 24 SNPs within the gene. No SNPs were found within the intron. Haplotype network analysis revealed higher diversity in the isolates collected on the resistant hybrids, compared to the susceptible hybrids (Figure 4a). The sequence diversity (pi) was 10X higher in the isolates originating from the resistant hybrids (pi=0.0037) compared to isolates coming from susceptible hybrids (pi = 0.00030) in Switzerland. Translation of the two exons identified 12 different amino acid sequences, with most of the variation in the amino acid sequence resulting in virulence at 40 dpi on Hybrid_R2 (Figure 4b).

**Figure 4.**
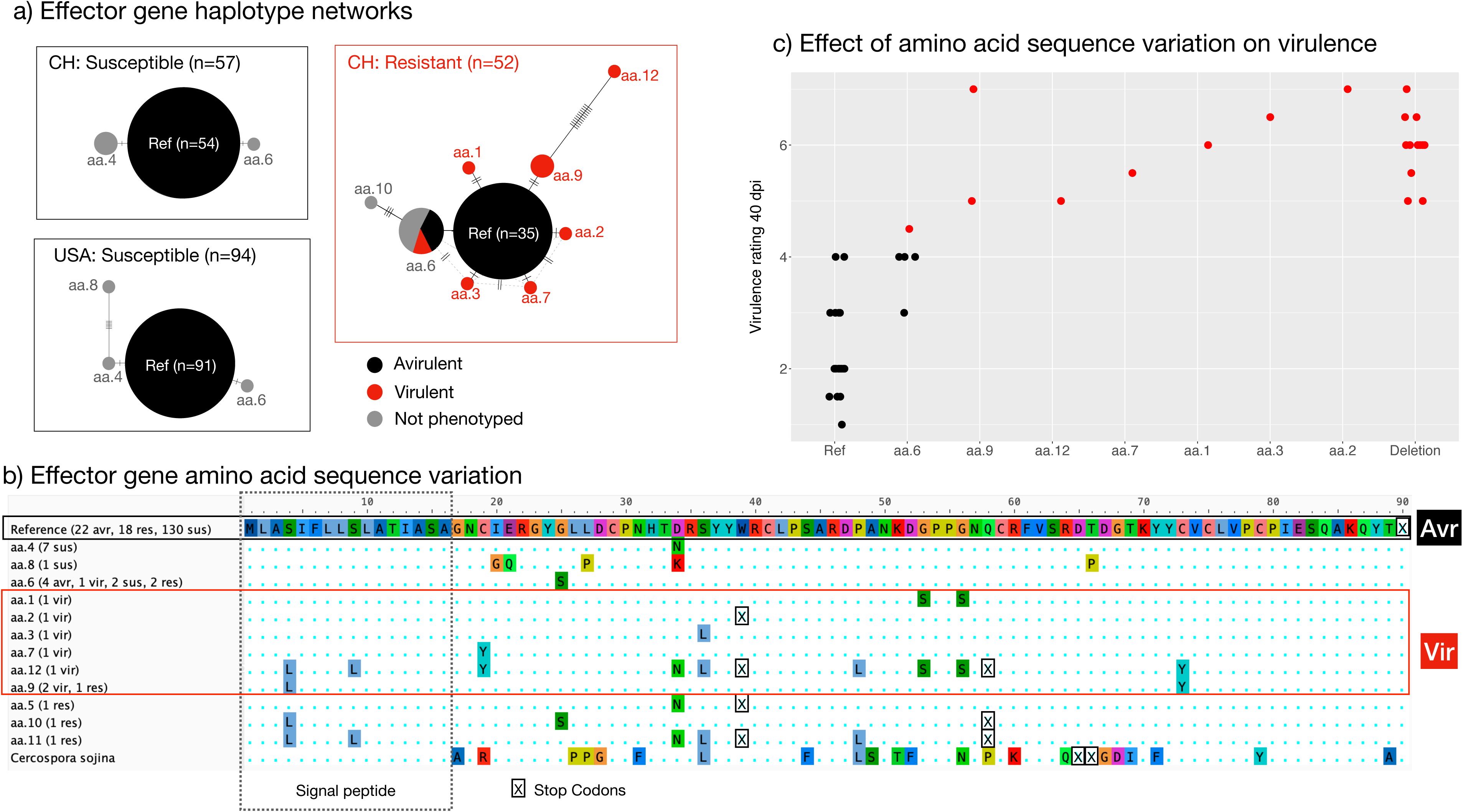
DNA and amino acid (aa) sequence variation in the candidate effector gene. a) Haplotype networks for isolates collected in Switzerland from resistant and susceptible hosts and in Renville USA. Ref = Identical to reference genome, ‘aa’ numbers correspond to the translated effector sequence variation shown in (b). b) Shows the 13 (incl. Ref) different amino acid sequences and the aa sequence for *Cercospora sojina*. In parentheses are the numbers and their host or virulence status (if phenotyped): sus = collected on susceptible hybrid, res = collected on resistant hybrid, virulence rating at 40 dpi on Hybrid_R2 vir >4 virulent, avr ≤4 avirulent. c) Effect of amino acid sequence variation (seen in b) and gene deletion on virulence rating (40 dpi Hybrid_R2).

To assess a potential fitness cost associated with loss of function of the putative effector, we tested if the loss or inactivation of the effector influenced the virulence of an isolate infecting resistant and susceptible hybrids in the 2022 field trial. We grouped isolates into two categories: those encoding the reference wild type isoform of the putative effector, and those encoding either deletions or alternative effector isoforms. We found no difference in disease rating between isolates with the deleted isoform or with alternative isoforms (Table S7). Across phenotyped isolates, we found that dpi, hybrid and effector gene isoform explained variation in disease rating, with significant interactions between all pairs of factors but no three-way interaction (Table S8). Considering each hybrid and dpi separately, we found that the effector isoform had little effect on an isolate’s virulence rating when infecting the susceptible hybrids but had a significant effect on an isolate’s virulence when infecting the resistant hybrid carrying *BvCR4* (Figure 5). Thus, there was no evidence for a reduction in fitness in strains lacking *AvrCR4* or encoding alternative isoforms of AvrCR4. Rather it appears that strains carrying deletions or alternative isoforms exhibit higher virulence on the susceptible hybrids (Figure 5, p-values ranging from 0.02-0.09, See Table S8).

**Figure 5.**
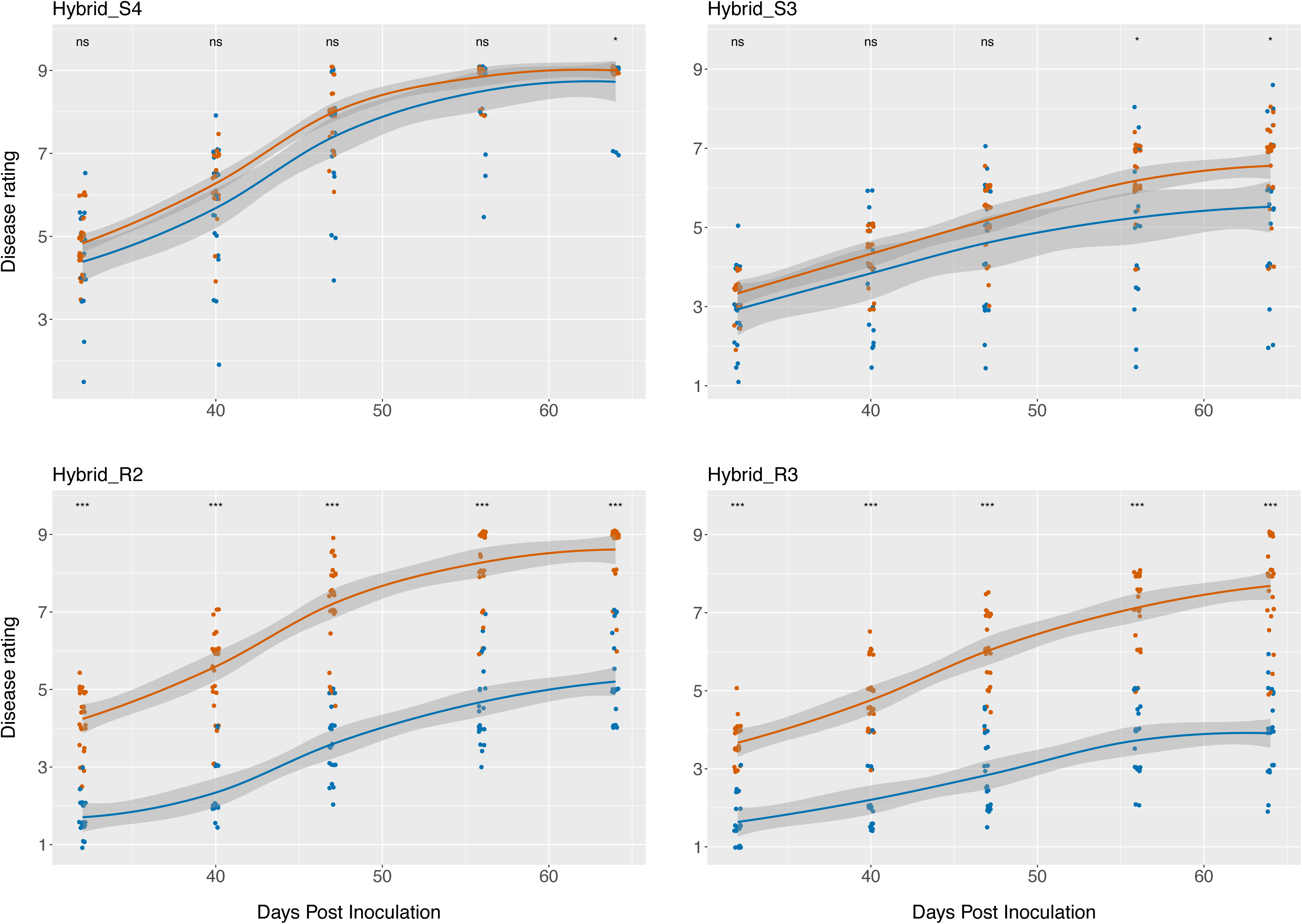
Field disease rating of isolates at five time points coloured according to the effector gene variant (blue=same as reference, orange=gene absent or sequence codes for different amino acids (inactivated)). Each panel shows the rating on one of four hybrids (2 susceptible and 2 resistant). Stars on top of the plots indicate significant differences between gene variants at each time point (ns=nonsignificant, *p<0.05, **p<0.01, *** p<0.001).

## Discussion

The identification and use of *BvCR4* represents an important advance in breeding for resistance to CLS, as it provides a high level of resistance to CLS without a reduction in sugar yield. Given the long history of boom-and-bust cycles associated with the release of major-effect R-genes, there was concern that *C. beticola* could evolve virulence against resistant hybrids carrying *BvCR4*, the first reported major R-gene deployed against this pathogen.

The first evidence that virulent isolates could evolve was observed as clusters of CLS-infected plants, hereafter called hotspots, in experimental fields planted with *BvCR4* hybrids in Switzerland in 2019. Although these hotspots did not emerge until late in the growing season, their occurrence suggested selection and local spread of *C. beticola* clones that were virulent to *BvCR4*. The discovery of these hotspots triggered our study to assess the emergence and spread of virulent strains in *C. beticola*.

Our GWAS used collections of *C. beticola* from plots planted with resistant *BvCR4* hybrids and susceptible hybrids lacking *BvCR4* growing in the same field. A subset of these *C. beticola* isolates were phenotyped under field conditions to test their virulence. The quantitative GWAS using 52 phenotyped isolates identified a region on Chromosome 1 associated with virulence, but the LOD scores were not significant. To increase the power to detect genes associated with virulence on *BvCR4*, we assumed that any allelic variants encoding virulence to *BvCR4* would be present at a very low frequency (e.g., at mutation-drift equilibrium) prior to the deployment of *BvCR4* hybrids. Hence, we scored virulence as a binary trait and included an additional 38 *C. beticola* isolates from susceptible hybrids growing in the same field scored as avirulent, although we lacked phenotypes for these strains. This binary phenotype approach identified the same Chromosome 1 region, with a significant LOD score of 9.3. A third GWAS based on k-mers, including only the phenotyped strains, identified the same Chromosome 1 region, with a very high LOD score of 21.3, as well as small intervals on Chromosomes 2 (LOD = 14.5) and 7 (LOD = 11.4).

A single predicted gene was found within the Chromosome 1 interval that was identified in all three GWAS analyses. This gene shares many properties found in functionally validated avirulence effectors in other plant pathogenic fungi, including: a) it encodes a small, secreted, cysteine-rich protein; b) it displays a presence-absence polymorphism and the absence allele associated with a gain in virulence occurs at a higher frequency on *BvCR4* hybrids compared to susceptible hybrids; c) its exons encode non-synonymous substitutions associated with virulence, but nucleotide diversity is lower in the introns and the signal peptide sequence; d) it is located in a genomic region that is rich in transposable elements and associated with RIP. Based on these properties, we named this candidate avirulence effector gene *AvrCR4*.

The main difference between *AvrCR4* and other cloned fungal avirulence effectors is that the overall level of DNA and amino acid diversity found in this study is lower than what has been reported for other avirulence effectors (Brunner et al., 2007; Brunner and McDonald, 2018; Dodds et al., 2006; Schürch et al., 2004). This lower diversity may reflect the very short period over which selection has been operating on this gene. In Germany, the first commercial cultivation of *BvCR4* sugar beet hybrids occurred in 2021 and in Switzerland in 2022. Based on the patterns of diversity we observe in this gene and comparisons to other well characterized avirulence effectors, we believe we have captured AvrCR4 at the very onset of its evolution. Firstly, the same wild-type *AvrCR4* sequence is found at a high frequency on both resistant and susceptible sugar beets growing on different continents (Figure 4a). In contrast, in *Z. tritici* the *AvrStb6* gene is present in every isolate analyzed to date and there is no predominant wild-type, despite the use of *Stb6* in European wheat cultivars for at least 100 (Dutta et al., 2021; Stephens et al., 2021). Secondly, there was little nucleotide diversity in *AvrCR4* in *C. beticola*. Amongst 224 isolates with AvrCR4 present, only 29 strains carried *AvrCR4* alleles that differed from the predominating wild type sequence. Among those 29 strains, we identified 11 distinct DNA haplotypes carrying 24 polymorphic sites encoding 13 AvrCR4 protein isoforms. No variation in the intron or signal peptide was observed. In comparison, in a global sample of 383 *Z. tritici* strains, 52 haplotypes encoding 44 AvrStb6 protein isoforms were found, with several polymorphisms found in the signal peptide (Stephens et al., 2021). In another study, evidence of diversifying selection acting on codons in AvrStb6 was found, suggesting that diversity in this gene has been maintained via a co-evolutionary arms between host and pathogen (Brunner and McDonald, 2018).

Finally, the *AvrCR4* presence/absence frequency is closely associated with the deployment of *BvCR4*. *AvrCR4* was present in nearly all *C. beticola* isolates sampled from locations where *BvCR4* has not yet been deployed (USA: 94/95). In experimental fields planted to both, hybrids carrying and lacking *BvCR4*, it was present in all isolates collected from hybrids lacking *BvCR4* and was often missing in isolates sampled from hybrids carrying *BvCR4* (33/89).

We believe that *AvrCR4* is the first avirulence effector to be discovered at the very onset of its evolution. This provides a unique opportunity to monitor in real time the changes in effector amino acid sequence that enable an escape from recognition by the cognate NBS-LRR protein. We also consider this to be a unique opportunity to search for compensatory mutations that could ameliorate any reductions in pathogen fitness caused by changing the function of the encoded wild-type avirulence effector protein.

Our results also provide important insights into how *C. beticola* can overcome resistant cultivars and if there are potential fitness costs associated with virulence. Standing genetic variation in *C. beticola* populations is the most likely source for the *AvrCR4* mutations creating virulence, as the gene deletion was detected in populations not exposed to *BvCR4* hybrids. Recent work has shown that populations of *Z. tritici* can generate and maintain vast reservoirs of functional genetic diversity facilitating rapid adaption to changes in their local environments (McDonald et al., 2022). Like *Z. tritici*, *C. beticola* has a mixed reproduction system and high gene flow (Rangel et al., 2020; Vaghefi et al., 2017), suggesting that its populations will also maintain high levels of standing genetic variation. Approximately half of the isolates collected from *BvCR4* hybrids were avirulent on *BvCR4* hybrids in our field phenotyping, illustrating that *BvCR4* hybrids can be colonized by *C. beticola* strains that are not specifically adapted to *BvCR4*. We speculate that other abiotic or biotic stresses could play a role. Although we did not test this specifically, we observed several instances of plants infected by *Rhizoctonia solani* in Rudolfingen in 2020 that might have compromised CLS resistance. Furthermore, experiments with mixed infections in *Z. tritici* showed that virulent strains were able to suppress the immune system of plants carrying the *Stb6* R-gene, enabling avirulent strains to infect the resistant plants (Bernasconi et al., 2023). We hypothesize that similar mechanisms could be operating in the CLS pathosystem. Finally, our study found little evidence of a fitness cost associated with the deletion or loss of function of AvrCR4. Strains carrying deletions for *AvrCR4* showed a similar disease development on sugar beet hybrids lacking *BvCR4*. This is in line with expectations of functional redundancy in effectors (Rouxel and Balesdent, 2017) and with previous work in *Z. tritici* (Zhan and McDonald, 2013). While we did not detect any fitness costs in the field environment, we are currently conducting additional lab-based experiments to search for fitness costs associated with loss of the *AvrCR4* function.

The identification of *AvrCR4* will benefit CLS management by enabling regular monitoring of the frequency of virulent mutants in CLS populations. Monitoring can support region-specific variety choice and management strategies aimed at reducing the overall CLS population and the frequency of virulent strains. These management strategies will likely utilize an integrated plant protection approach that combines precise timing of fungicide applications with agronomic practices such as crop rotation and tillage to reduce the buildup of inoculum in the soil (Harveson et al., 2009; Khan et al., 2008; Khan and Khan, 2010; Secor et al., 2010) (Khan et al., 2008; Khan and Khan, 2010; *Jacobsen and Franc, 2007; Secor et al 2010*). Varieties that combine *BvCR4* with other sources of resistance can provide a strong CLS background resistance that will further contribute to a reduction of the CLS population and increase the durability of *BvCR4*. This latter approach was shown to be effective in oilseed rape, where the adaptation time of *L. maculans* to the resistance gene *Rlm6* was doubled by protecting this resistance with polygenic background resistance (Brun et al., 2010; Delourme et al., 2014; Pilet-Nayel et al., 2017).

### Experimental Procedures

#### Collections of leaves infected by CLS

In 2019 a field in Switzerland (Table S9) was planted with two experimental sugar beet hybrids; Hybrid_R1 carrying *BvCR4* in a genetic background with low quantitative resistance, and Hybrid_S1 carrying only a low level of quantitative resistance. Both hybrids were examined in 2019 for evidence of CLS infections. Infected leaves were collected from hotspots exhibiting CLS symptoms and stored for later isolation.

In 2020, four sugar beet experimental hybrids with different levels of resistance to CLS were planted at two field sites in Switzerland separated by 60 km (Table S9); Hybrid_R2 with *BvCr4* embedded in a background with low quantitative resistance, Hybrid_R3 with *BvCr4* embedded in a background with moderate quantitative resistance, Hybrid_S2 with only low quantitative resistance and Hybrid_S3 with moderate quantitative resistance (Table S1). In Hendschiken, 12 rows of each hybrid were planted in adjacent plots and in Rudolfingen, 18 rows of each hybrid were planted in adjacent plots. Sugar beets were sown in March and harvested in early November. The same four experimental hybrids were planted in similar experimental plots at two field sites in Germany (Table S1 and S9). No fungicides were applied at any of the field sites.

We used two sampling strategies, one temporal (Swiss sites only) and the other spatial (Swiss and German sites), to obtain leaves infected by *C. beticola* (Figure S1). The temporal sampling strategy was designed to identify infected plants in each plot and monitor how CLS infection spreads from a solitary infected plant to neighboring plants to form a cluster of CLS-infected plants, called a hotspot. We identified every newly infected plant in each plot at weekly intervals from July 21^st^ to September 24^th^ 2020. Every infected plant was given an alphanumeric code and its position was marked with a labeled flag to avoid resampling. One young leaf with CLS lesions was taken from each infected plant. The excised leaves were transported back to the laboratory for *C. beticola* isolation. A GPS location was recorded for the first infected plant in each hotspot. A maximum of five plants was sampled from each hotspot.

Transect sampling was conducted in all plots in Switzerland and Germany at the end of the 2020 season (13^th^ and 15^th^ October in Switzerland and 17^th^ September in Germany). A total of 7 transect lines each 3 m apart were made for each transect site, giving a total of 21 leaves per transect site (see Figure S1). Two transect sites were sampled for each hybrid, yielding 42 leaf samples total for each hybrid in each plot at both sites in Switzerland and Germany.

### Pathogen isolation and DNA extraction

Leaf pieces bearing CLS lesions were cut from each leaf and surface-sterilized by placing them in a 2% sodium hypochlorite solution for 3 min and then rinsing them twice in sterile water. The leaf pieces were then placed in glass petri dishes on a sterile plastic screen net on top of a piece of moist filter paper. The petri dishes were incubated at 24°C for 48-72 hours to induce sporulation. Spores from single sporulating lesions were gathered using hand-made fine glass needles and transferred to fresh potato dextrose agar (PDA, Difco). The plates were cultured under constant darkness at 24°C for 7-10 days. Any plates contaminated by other fungi were discarded.

To obtain fresh mycelia for DNA extraction, we transferred hyphae from pure culture plates to 100 mL flasks with 50 mL potato dextrose broth (PDB, Difco). DNA for 103 isolates was extracted with a Kingfisher Flex robot (Thermo Fisher Scientific, U.S.A.) following the recommended protocol for fungal samples (Fungus Sbeadex, LGC Genomics GmbH) and DNA for the remaining 153 isolates was extracted using DNeasy Plant Mini Kits (Qiagen) with a modified protocol; 3.5 μL of proteinase K was added to each sample at the lysis step. DNA was quantified using the Qubit 2.0 Fluorometer (Themo Fisher Scientific, U.S.A.) with a Qubit dsDNA HS Assay kit (Thermo Fisher Scientific, U.S.A.) and Nanodrop (ThermoFisher Scientific, U.S.A.). Species identity was confirmed using PCR (Knight and Pethybridge, 2020).

For the reference genome, *C. beticola* strain “Hevensen” was grown in liquid media by transferring mycelium from a V8 agar petri dish (500 ml of V8 juice, 500 ml distilled water, 3 g of CaCO_3_ and 15 g of agar) to 225 ml of CB medium (1 g/l casein, 1 g/l yeast extract, 10 g/l D-Glucose, 10 mM NaCl, 0.02 mM FeCl_3_x6H_2_O, 0.1 mM ZnSO_4_x7H_2_O, 7.9 µM CuSO_4_x5H_2_O, 0.4 mM MnSO_4_xH_2_O, 0.9 mM H_3_BO_3_, 4.2 mM Ca(NO_3_)_2_x4H_2_O, 1.5 mM KH_2_PO_4_, 1 mM MgSO_4_x7H_2_O). The culture was incubated for 6 to 8 days at 24°C at 100 rpm. Mycelium was ground to a fine powder in liquid nitrogen. DNA was extracted using the method described by (Huang et al., 2018) with minor modifications. DNA was repaired and end-prepped using NEBNext FFPE DNA repair mix (NEB) and the NEBNext Ultra II End Repair/dA-Tailing Module (NEB), respectively. Small DNA fragments were removed from each sample using a Short Read Eliminator reagent (Circulomics).

### Genome data acquisition and assembly

The Hevensen reference isolate was prepped for Oxford Nanopore Technology (ONT) sequencing using SQK-LSK109 library kit (ONT). Each library was loaded onto three FLO-FLG001 (R9.4.1 pore) flongle cells (ONT) and run for 24 hours on MinION or GridIONx5 instruments (ONT). Raw data was base called using Guppy (version 4.0.15, default parameters) and assembled with miniasm (version r179; default parameters). The resulting draft assembly was polished with Racon (version 1.4.13, input long reads, default parameters), Medaka (version 1.2.0, input long reads, default parameters) and Pilon (version 1.23, input short reads, default parameters). Polished sequences were size-ordered and named in consecutive order as chromosomes 1 through 11. Two short contigs were annotated via Uniprot Blast and contained rDNA (CB_CbHevensen-1_hq-v2.rDNA) and mtDNA (CB_CbHevensen-1_hq-v2.mitochondrial). Final sequences were structurally annotated with Funannotate (version 1.8.3, RNA-seq evidence – NCBI PRJNA294383, protein evidence - NCBI RefSeq Fungi 2021-05-06, otherwise default fungal parameters) and CodingQuarry-PM (run_CQ-PM_mine.sh, version 2.0) (Table S6a). RepeatMakser (v4.0.6) and RepeatModeler (v1.0.11) were used to identify repetitive elements in the genome (- species Fungi) (Figure S3). Signatures of RIP across the genome were investigated using RIPPER (van Wyk et al., 2019) (Table S6b).

DNA from the 256 field-collected isolates was sequenced using Illumina paired end (150 bp) sequencing (sequence provider: Novogene). In addition, raw reads from 199 isolates were downloaded from the SRA (https://www.ncbi.nlm.nih.gov/sra) using SRAtoolkit (v2.0) for downstream analyses. We performed multiple analyses to check the quality of the isolate sequence data, see Supplementary Information (SI) for details. Briefly, we used k-mer based approaches to identify potential contaminants or mixed cultures by examining the heterozygosity and genome coverage. We estimated heterozygosity and sequence coverage using Smudgeplot (https://github.com/KamilSJaron/smudgeplot) and removed isolates with high heterozygosity (>0.1%) or lower than expected genome coverage. We drew a k-mer based distance tree with mashtree (Katz et al., 2019) and removed isolates that were genetically distinct and potentially different species. Sequence data processing and variant discovery are described in detail at https://github.com/jessstapley/CercosporaGWAS. After trimming reads were mapped to the Hevensen reference genome using bwa-mem (v0.7.17). Samples that did not map well to the reference genome (< 80% properly paired reads) were removed. After mapping, quality checking and removal of sequence duplicates, we retained 422 isolate genomes (232 sequenced by us, 190 from SRA) for further analyses. See SI.1 and SI.2 for details and Table S1 for information about each isolate, including why they were removed from further analyses.

Variant calling was done using the GATK (v4.2.4.1) Germline Short Variant Discovery pipeline, following their Best Practices recommendations (https://gatk.broadinstitute.org/hc/en-us/sections/360007226651-Best-Practices-Workflows). For the GWAS analyses we used only biallelic SNPs and we removed markers with minor allele frequency <0.05. The SNP error rate based on isolates sequenced twice (sequence duplicates, n=29) was 0.00125. Clones were identified using an IBS distance matrix from LD pruned data (13,696 SNPS) (See SI.3 for details). Following clone correction, we retained 163 unique isolates from Switzerland and Germany, and 95 unique isolates from the U.S.

### Field phenotyping of virulence for strains used in the GWAS

Sixty fungal inocula were tested in total. Fifty-two unique genotypes originating from Swiss plots were tested, 49 from plots planted to *BvCR4* hybrids (18 isolates from Hybrid_R2, 31 isolates from Hybrid_R3) and three from the susceptible hybrids (2 from Hybrid_S3, 1 from Hybrid_S4). One putatively virulent clone that was widespread in one of the experimental fields was replicated five times, giving a total of 56 field-collected isolates. Four additional reference inocula were used as controls with known virulence properties but were not sequenced nor included in the GWAS analyses.

Inoculum of the *C. beticola* strains was produced at KWS in Einbeck, Germany by first growing on V8 agar (see S1.4 for details). Next, groups of 30, 12-week old plants of Hybrid_R2, were inoculated with each of the isolates used in the GWAS by spraying suspensions of ∼25,000 hyphal fragments/spores per ml onto each group of plants until runoff. Inoculated plants were then covered with plastic sheets draped over metal frames and incubated for 5 days without supplemental lighting at 25°C and kept at 100% RH. The first symptoms were observed at 14 dpi for most isolates. Inoculum for each isolate was harvested between 28 and 42 dpi by removing leaves that were fully covered with CLS lesions. Leaves were dried, milled and mixed with semolina flour.

Virulence assessments were conducted in 2022 on a field site in Germany. To minimize the risk of infection by local strains of *C. beticola*, the field had not been planted to sugar beet for about 20 years (corn-wheat rotation), the nearest sugar beet field was 800 m distant, and the experimental field site was surrounded by a 50 m border of maize. Seeds treated with the fungicide Tachigaren 70WP (Hymexazole) and the insecticide Force 20CS (Tefluthrine) from four sugar beet hybrids with varying levels of CLS resistance (Table 1) were planted on 19^th^ April 2022 in a randomized block design with 6-row plots that were 0.45 m wide and 5 m long. Each isolate was inoculated onto a block of plots that contained all 4 hybrids. Weeds were controlled with three herbicide applications and aphids were controlled with two insecticide applications during the growing season.

Plots were inoculated after row closure on 27^th^ July 2022. Inoculation was performed by spreading 60 g of leaf inoculum mix (10 g milled leaf inoculum + 50 g semolina flour) onto one row (3^rd^ or 4^th^ row, 5 m length, approx. 25 plants) of each 6-row plot. Inoculation rows were sprayed with water containing an adjuvant (Silwet 0.05%) prior to inoculation so that most of the inoculum mix would stick to the leaf canopy. Following the inoculation, all plots were watered twice a week for 6 weeks to facilitate infection and disease progress.

Approximately 4 l water/m^2^ was applied each week with a tractor-mounted crop protection sprayer. We recorded the disease rating at 32, 40, 47, 56 and 64 dpi. We analysed changes in disease rating using linear models and least significant differences to make post hoc comparisons (Bonferroni-adjusted) between mean rating at each time point. Disease rating was bimodal at 40 dpi on the resistant hybrid Hybrid_R2 (Figure S4), providing a clear distinction between virulent (rating >4 on KWS rating scale) and avirulent (rating ≤ 4) strains and enabling us to code isolates as either virulent or avirulent for some of the GWAS analyses.

### GWAS analyses

SNP-based GWAS were performed using GAPIT (http://zzlab.net/GAPIT). We performed two SNP-GWAS analyses using different phenotype data as described above. We used a Mixed Linear Model (MLM), which accounts for relatedness of individuals, and included three principal components into the model as covariates (random effects). The model fit was assessed by inspecting the Q-Q plots (Figure S5). We repeated the analysis using quantitative virulence ratings at other time points (Figure S6). A quantitative GWAS using the Hybrid_R2 virulence rating at 47 dpi identified the same peak that was found using the rating at 40 dpi.

We conducted a k-mer GWAS (see https://github.com/jessstapley/CercosporaGWAS for details) as a recent analysis conducted in *Z. tritici* (Dutta et al., 2022) showed that these can provide more power to detect associations. We used recommendations and scripts from https://github.com/voichek/kmersGWAS, but used GAPIT to run the GWAS. In brief, KMC was used to count k-mers with a length of 31 bp. The k-mer plink output file (.bed) was imported into plink (v1.9, https://www.cog-genomics.org/plink/) and converted to a vcf file (--recode vcf). The vcf file was split into 99 smaller files (containing 20,000 k-mers each), then converted to a hapmap file using R::vcfR (https://github.com/knausb/vcfR). GAPIT GWAS was then run on each of these smaller files. The k-mers were mapped to the reference genome using bowtie2 (v2.3) and the results were concatenated into chromosome files for plotting. The k-mer GWAS was performed using only 47 of the phenotyped strains because we could not count k-mers in all 52 strains.

### Annotation of the GWAS interval

Genes within the GWAS interval were identified using the reference genome annotation file (gff3). The function of the putative effector was confirmed using EffectorP (v3.0) (https://effectorp.csiro.au/ (Sperschneider and Dodds, 2022)). Additional annotation of the GWAS interval was performed to identify possible transposable elements (TEs) using Repeat Masker (v4.0.6) and One Code to Find them all (Bailly-Bechet et al., 2014).

SNP genotypes across the GWAS interval were obtained for every isolate using bcftools (samtools v1.9). Individuals with missing genotype calls spanning the entire candidate gene were recorded as ‘gene deleted’ for the candidate gene. Individuals with SNP genotype calls within the gene were recorded as ‘gene present’. We compared the frequency of gene presence or absence between resistant and susceptible hybrids in Germany and Switzerland using a binomial exact test.

## Acknowledgments

We are grateful to Dr. Julien Alassimore, Sophie Kuhn and Vivienne Hanke for support in the lab.

## Data availability statement

All sequence data has been deposited on the NCBI SRA (SRA NUMBER). The genome is available at DOI. All other data is available in the Supplementary Information.

