## Supplementary material for "Genome-wide association studies reveal a rapidly evolving candidate avirulence effector in the Cercospora leaf spot pathogen *Cercospora beticola*": Supplmentary Information

### Supplementary Information (SI): Methods

#### SI.1. Filtering sequence data

We used k-mer based approaches to identify potential contaminants or mixed cultures. To verify that isolations made in the lab were composed of a single clone, we first counted k-mers with KMC (https://github.com/refresh-bio/KMC) (Kokot et al., 2017) and then estimated heterozygosity and sequence coverage using Smudgeplot (https://github.com/KamilSJaron/smudgeplot). Four isolates had high heterozygosity (>0.1) and five had a k-mer based genome coverage lower than the expected coverage based on the number of reads (number of reads * read length)/haploid genome size < 0.5). Next, we investigated genetic similarity to identify isolates that were genetically distant, representing potentially different species, subspecies or contaminated cultures. We used the program mashtree (Katz et al., 2019) to draw a k-mer-based neighbor joining tree. Thirteen isolates that were highly divergent were excluded from further analyses.

#### SI.2. Variant Calling and Filtering

Variant calling was done using the GATK (v4.2.4.1) Germline Short Variant Discovery pipeline, following their Best Practices recommendations (https://gatk.broadinstitute.org/hc/en-us/sections/360007226651-Best-Practices-Workflows). GATK default filters are: MQ=20.0, ReadPosRankSum_lower=-2.0, ReadPosRankSum_upper=2.0, MQRankSum_lower=-2.0, MQRankSum_upper=2.0, BaseQRankSum_lower=-2.0, BaseQRankSum_upper=2.0. In addition to the GATK default filters we excluded rDNA and mtDNA. We also removed indels and loci with >50% missing genotypes, as well as loci that did not meet the following criteria: depth <20000, mapping quality (MQ) >50, quality depth (QD) >2.0, Symmetric Odds Ratio (SOR) <3, and Phred-scaled p-value using Fisher's exact test to detect strand bias (FS) <60. For the GWAS analyses we used only biallelic SNPS and we removed markers with minor allele frequency <0.05. Principal component analysis was performed on LD-pruned SNP datasets (see https://github.com/jessstapley/CercosporaGWAS for details). The SNP error rate based on isolates sequenced twice (sequence duplicates, n=29) was 0.00125.

**SI.3. Clone Correction**

Clones were identified using an IBS distance matrix from LD pruned data (13,696 SNPS). To determine the appropriate genetic similarity threshold, we used SNP data from the known sequence replicates. We calculated the genetic similarity of SNP data known to come from the same genotype and found it ranged between 0.9968-0.9990. We considered three thresholds (0.990, 0.995, 0.999) to identify clones. A threshold of 0.999 was too strict, with only one pair of known sequence replicates correctly recorded as having the same genotype. The thresholds of 0.990 and 0.995 gave identical results and correctly assigned all sequence replicates as the same genotype. Thus, isolates with a genetic similarity >0.990 were considered a single genotype or clone. Following clone correction, we retained 163 unique isolates from Switzerland and Germany, and 95 unique isolates from the publicly available genomes of isolates coming from the USA.

**S1.4. Production of Inoculum**

Inoculum of the *C. beticola* strains was multiplied at KWS in Einbeck, Germany between March and June 2022. The strains were cultured on Petri plates containing V8 agar. A 5 mm plug of mycelium for each isolate was transferred onto a petri plate and incubated at 20°C under 12-hour darkness: 12-hour fluorescent light until the plates were at least 50%

colonized, requiring approx. two weeks. Mycelium plugs from these starter plates were then transferred onto fresh V8 plates (3 plates per isolate) and incubated as described above for another 2 weeks. Fully colonized plates were blended in 50 ml of distilled water to make a suspension. Approximately 5 ml of this suspension was poured into 25 new V8 agar plates, which were then rotated to evenly spread the liquid on the agar surface. The plates were then placed into a cardboard box and incubated in the dark at 20°C for 2 weeks. Inoculum consisting of mycelial fragments and spores was harvested by adding 5 ml of distilled water per plate followed by gently scraping off the fungal cells growing on the agar surface. The resulting fungal mass, containing both hyphal fragments and spores, was filtered through cheesecloth and then adjusted to a concentration of 25,000 hyphal fragments/spores per ml. The suspension from each isolate was amended with 0.1% Tween 20 (Sigma-Aldrich) prior to inoculation.

Hybrid_R2, an experimental hybrid shown in earlier experiments to be susceptible to *C. beticola* under greenhouse conditions, was planted in trays filled with peat soil mix (HAWITA). After 2 weeks, seedlings were transplanted into new trays and kept at 16°C for 6 weeks under diffuse light. Plants were then transferred into plastic pots (11x11x10 cm) filled with peat soil mix and arranged in groups of 30 plants in a greenhouse chamber set at 20°C with 16h fluorescent light per day for four weeks. Plants were fertilized weekly with 0.8% Kamasol green (10N-4P-7K). The distance between each group of 30 plants was 0.5 m to minimize the potential for cross-contamination of the inoculum produced for each strain.

Each group of thirty 12-week-old plants was inoculated with one of the 56 isolates used in the GWAS. Suspensions of spores and hyphal fragments were sprayed onto each group of plants until runoff with a hand-held atomizer. Inoculated plants were then covered with plastic sheets draped over metal frames and incubated for 5 days without supplemental lighting at 25°C and kept at 100% relative humidity (RH). After the incubation, plants were kept under a 16 h photoperiod at 25°C/20°C (day/night) with 60-80% RH. The first symptoms were observed at around 14 dpi for most isolates. Inoculum for each isolate was harvested between 28 and 42 dpi by removing leaves that were fully covered with CLS lesions. The removed leaves were placed in paper envelopes and dried at 35°C for 3 days. The dried leaves were then milled in a kitchen blender for 30 sec, thoroughly mixed with semolina flour (40 g of milled leaf inoculum added to 200 g of flour) and then transferred into labelled plastic boxes.
